# Profiling host-pathogen dynamic in a 3D human epidermal infection model and real-time sensing of IL-1α as a tissue-damage biomarker

**DOI:** 10.64898/2026.11.19.767804

**Authors:** Liu Yang, David Valero Calvo, Sihui Xu, Mafalda Moura, Kuo Liao, Alexander Zink, Aimo Kannt, Nicole Ziegler, Maurice Michel, Cédric Badiou, Alberto J. Martín-Rodríguez, Morteza Hasanpouramiri, Onur Parlak

## Abstract

Skin infections begin with superficial colonization and progress to host tissue damage, yet the models used to study this process and the sensors intended to monitor it are rarely developed together. Here we combine a standardized three-dimensional human epidermal equivalent (HEE) infection model with an electrochemical immunosensor. By using a surface-restricted inoculation method, this model supported live pathogenic bacterial exposure on intact epidermis for up to seven days without bacterial contamination of the basolateral culture medium. Over the 7-day course, while commensal *Staphylococcus epidermidis* remained confined to the stratum corneum with preserved tissue integrity, *Pseudomonas aeruginosa* and methicillin-resistant *Staphylococcus aureus* (MRSA) caused progressive epidermal barrier damage and an inflammatory signaling cascade. Longitudinal cytokine profiling revealed distinct host-response profiles, identifying interleukin-1α (IL-1α) as the most consistent and responsive indicator of the transition from colonization to active infection across both *P. aeruginosa* and MRSA groups. A laser-patterned gold/polyimide electrode functionalized with anti-IL-1α antibodies resolved the rise in IL-1α from bacterial colonization (day 4) to severe infection (day 7) by electrochemical impedance spectroscopy, providing state discrimination rather than quantitative equivalence with ELISA. Overall, this integrated approach establishes a robust, animal-free platform that links infection-driven epidermal damage with biomarker-based electrochemical detection, providing a foundation for the development of next-generation *in vitro* diagnostics.

## 1. INTRODUCTION

Skin and soft tissue infections (SSTIs) represent a major global health burden, a challenge further compounded by the escalating crisis of antimicrobial resistance (AMR)^[1,2]^. In 2019, bacterial AMR was estimated to be directly attributable to 1.27 million deaths and associated with 4.95 million deaths worldwide^[3]^. However, this resistance crisis represents only the clinical endpoint of a complex biological process.

At the biological level, the pathogenesis of many SSTIs begins with microbial colonization of the skin surface^[4]^. Under homeostatic conditions, the skin supports a diverse microbiota that coexists with the host and contributes to cutaneous homeostasis. The intact epidermal barrier plays a central role in maintaining this equilibrium by limiting microbial invasion while preventing excessive host immune activation^[5,6]^. However, upon barrier compromise induced by, for example, physical trauma, altered permeability, or underlying dermatopathology, some opportunistic pathogens (e.g., *Pseudomonas aeruginosa* and *Staphylococcus aureus*) can exploit the microenvironmental breach to colonize and proliferate. Under favorable conditions, these microorganisms can also form biofilms, which can enhance bacterial tolerance to host immune defenses and antimicrobial treatments^[7]^. During this spatiotemporal transition from subclinical colonization to acute infection, keratinocytes act as crucial epithelial sentinels that bridge the physical barrier and the underlying immune system. Upon structural disruption and exposure to bacterial virulence factors, keratinocytes release damage-associated molecular patterns (DAMPs), which activate signaling cascades that partially overlap with those triggered by pathogen-associated molecular pattern (PAMP) recognition, thereby mobilizing localized host defenses^[8,9]^. Resolving the dynamic profiles of these signaling molecules is therefore necessary for defining when microbial exposure has progressed to epidermal damage.

Despite extensive research, conventional biological models for deciphering these host-pathogen interactions exhibit significant limitations^[10]^. *In vivo* animal models, predominantly rodents, are constrained by pronounced interspecies physiological discrepancies in skin architecture and immunology^[11]^, alongside increasing ethical and regulatory emphasis on the principles of Replacement, Reduction, and Refinement (3Rs)^[12]^. *Ex vivo* human skin models, while offering excellent physiological relevance, are hindered by donor scarcity, high inter-donor variability, and low scalability^[13]^. Furthermore, conventional two-dimensional (2D) cell cultures lack the essential three-dimensional (3D) architecture required to reproduce epidermal barrier function. Although direct co-culture of 2D cells with live bacteria is possible, prolonged co-culture can lead to rapid bacterial overgrowth, rapid bacterial overgrowth and premature host-cell damage, making it challenging to reproduce the spatial and temporal dynamics of continuous bacterial colonization under controlled conditions^[14]^. These methodological gaps highlight the necessity for a scalable *in vitro* human system that preserves 3D barrier architecture while sustaining live host-pathogen interactions over extended periods.

Concurrently, the rapid evolution of flexible bioelectronic systems and wearable electrochemical sensors has opened new opportunities for the real-time, multiplexed monitoring of physiological biomarkers^[15]^. Recent studies have demonstrated the potential of smart sensing platforms to monitor inflammatory and metabolic signatures directly from wound exudates and other biological fluids^[16,17]^. More broadly, a critical validation gap persists between sensor development under simplified laboratory conditions and performance evaluation in biologically complex environments, hindering the translation of biosensing technologies into clinically relevant applications. Many biosensors are initially optimized in buffer systems that do not adequately reproduce the effects of host proteins, tissue-derived components, cellular debris, or bacterial extracellular polymeric substances (EPS), all of which can contribute to biofouling and alter sensor performance^[18,19]^. Consequently, promising biosensing platforms often progress from simplified benchtop characterization directly to animal studies, limiting opportunities for controlled, reproducible, and high-throughput optimization. A physiologically relevant *in vitro* infection model could therefore provide an important intermediate testing environment in which biosensor performance can be evaluated under biologically meaningful conditions while reducing reliance on animal experiments during early-stage development.

To bridge the translational gap between biological modelling and bioelectronic engineering, this study presents an integrated ‘biology-to-device’ platform that combines a standardized 3D epidermal infection model with a customized rapid electrochemical sensing platform. As an alternative to early animal testing, we established a standardized 3D human epidermal infection model to serve as the testbed. By resolving the spatiotemporal dynamics of both commensal and opportunistic pathogens within this 3D architecture, we identified interleukin-1α (IL-1α) as a host-derived signal associated with epidermal barrier damage during opportunistic infection. To assess whether this infection-associated host response could be monitored using a rapid electrochemical approach, we further developed an antibody-functionalized flexible Au/polyimide electrochemical sensor for IL-1α detection and evaluated its ability to resolve conditioned media collected at different stages of infection in the model. Rather than aiming to replace quantitative ELISA, we demonstrate the potential of a flexible electrochemical platform to translate infection-associated host responses into rapid electrochemical readouts. Overall, this integrated approach provides a potentially animal-sparing in vitro framework for evaluating next-generation medical bioelectronic devices under pathophysiologically relevant conditions.

## 2. RESULTS

### 2.1. Barrier disruption amplifies acute bacterial colonization and epidermal damage

Establishing physiologically relevant *in vitro* infection models is technically demanding. Conventional direct co-culture of live bacteria with 3D skin equivalents typically results in basolateral culture medium contamination within hours^[20]^, potentially compromising costly tissue models that take weeks to generate. To overcome these limitations, we adapted the surface-restricted inoculation methodology established by Rikken et al.^[21]^. By using a confining cylinder and an air-drying step, bacterial colonization was restricted to the epidermal surface. Building upon this technique, we established an acute infection model using human epidermal equivalents (HEEs) to mimic opportunistic infections in which pathogens exploit pre-existing skin barrier defects (**Figure 1A**).

**Figure 1.**
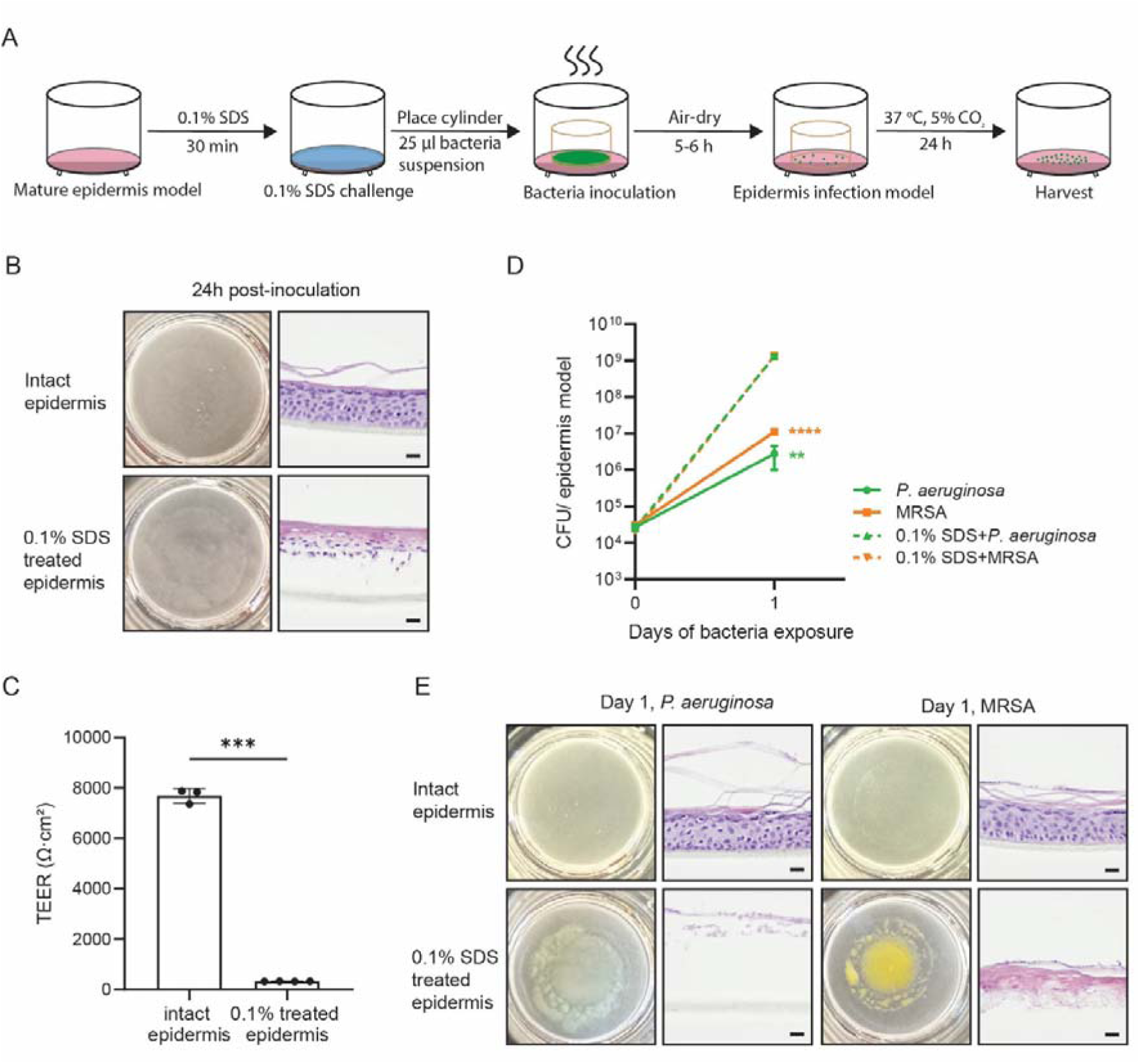
Establishment and evaluation of the acute bacterial infection model on 3D human epidermal equivalents (HEEs). (A) Schematic illustration of the experimental workflow adapted from the surface-restricted inoculation methodology. Mature HEEs were either kept intact or pre-challenged with 0.1% SDS for 30 min to disrupt the barrier. Bacterial suspensions (25 μL) were inoculated onto the surface using a confining cylinder, followed by air-drying and a 24-hour incubation at 37°C to prevent culture medium overgrowth. (B) Macroscopic top-down views and representative H&E histological cross-sections of the HEEs at 24 h post-inoculation in two groups (intact and 0.1% SDS-treated epidermis). (C) Transepithelial electrical resistance (TEER) measurements quantifying the severe loss of barrier integrity following the 0.1% SDS challenge (\*\*\**p* < 0.001, Student’s *t*-test). (D) Quantitative colony-forming unit (CFU) analysis showing amplification of bacterial burden (*P. aeruginosa* and MRSA) on SDS-compromised models compared to intact models at day 1 post-inoculation (\*\**p* < 0.01, \*\*\*\**p* < 0.0001, determined by Welch’s *t*-test on log-normally distributed data). (E) Macroscopic images and corresponding H&E-stained cross-sections of intact and SDS-treated epidermis infected with *P. aeruginosa* and MRSA for 24 hours. Scale bars in (B) and (E), 20 μm.

First, the structural integrity of the epidermis was compromised using a 0.1% SDS challenge prior to bacterial inoculation. Macroscopic observations and histological examination of hematoxylin and eosin (H&E)-stained cross-sections (Figure 1B) confirmed that the SDS treatment disrupted the superficial stratum corneum. This loss of barrier integrity was quantified by transepithelial electrical resistance (TEER) measurements. The intact epidermis exhibited a TEER of approximately 7800 Ω·cm², which sharply declined by over 95% following the SDS challenge (*p* < 0.001, Figure 1C), indicating a profound loss of barrier integrity.

Within this compromised microenvironment, bacterial proliferation was markedly enhanced. At 24 hours post-inoculation (day 1), colony-forming unit (CFU) analysis (Figure 1D) revealed that bacterial burden on the intact epidermis remained constrained, reaching approximately 10^6^ CFU/model for *P. aeruginosa* and approximately 10^7^ CFU/model for *MRSA*, whereas on the SDS-treated epidermis both species reached approximately 10^9^ CFU/model (*p* < 0.01 for *P. aeruginosa*, *p* < 0.0001 for *MRSA*). This corresponds to an increase of approximately 2.5–3 orders of magnitude for *P. aeruginosa* and approximately 2 orders of magnitude for *MRSA*. Histological evaluation at day 1 (Figure 1E) further corroborated these findings, demonstrating that the residual epidermal architecture in the SDS-treated models was rapidly and extensively degraded under this high bacterial load. In contrast, intact HEEs successfully maintained relative structural stability and limited the bacterial proliferation during this 24-hour acute phase.

### 2.2. Differential Spatiotemporal Dynamics of Long-Term Bacterial Colonization

While the acute model demonstrates early-stage barrier vulnerability, it does not capture the extended spatiotemporal evolution of infection. To investigate these long-term dynamics, we used a longitudinal infection model on intact HEEs cultured over a 7-day period (**Figure 2A**). To establish a physiological baseline of host-microbiome homeostasis, we included *S. epidermidis*, a prevalent skin commensal, as a biological control alongside the invasive pathogens. Monitoring revealed divergent colonization kinetics among the three evaluated bacterial strains. The commensal strain *S. epidermidis* established a stable homeostatic plateau at approximately 10^6^ CFU/model from day 2 to day 7 (Figure 2B). Accordingly, the epidermal models inoculated with *S. epidermidis* maintained structural integrity and exhibited 100% survival rate throughout the extended culture period (Figure 2C), remaining morphologically indistinguishable from the PBS-treated controls.

**Figure 2.**
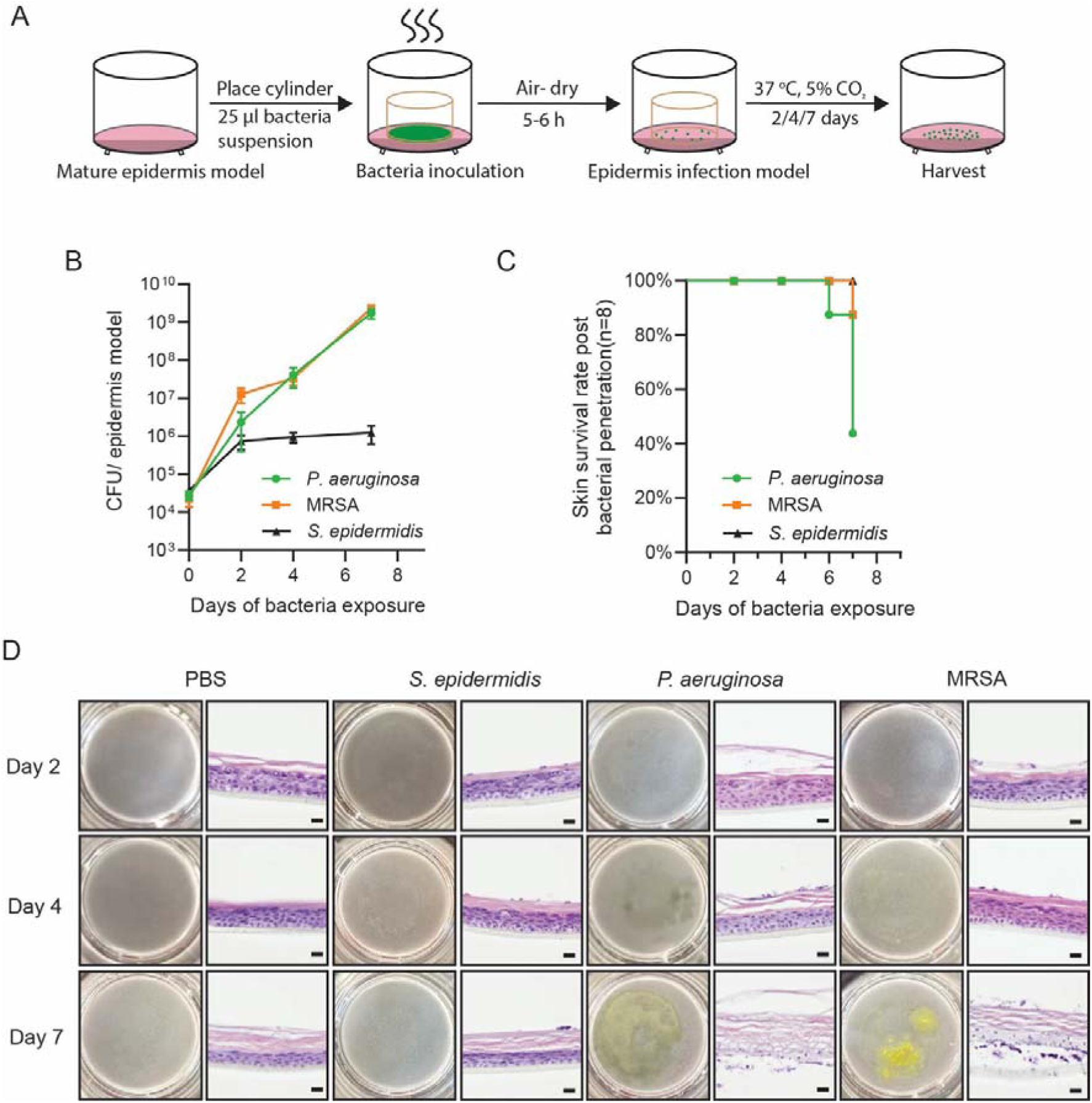
Spatiotemporal dynamics of long-term bacterial colonization on intact human epidermal equivalents. (A) Schematic illustration of the 7-day longitudinal infection model workflow. (B) Longitudinal quantification of bacterial burden (CFU/epidermis model) at day 0, 2, 4, and 7, highlighting the unchecked proliferation of pathogens versus the stable colonization of the commensal strain. Data are presented as mean ± SD. Statistical significance was determined by two-way ANOVA with Dunnett’s multiple comparisons test on log-transformed data, using S. epidermidis as the control reference (n = 3–6 per group, *p < 0.05, ****p < 0.0001). (C) Kaplan-Meier survival curves of the epidermal models following exposure to *P. aeruginosa*, MRSA, and the commensal *S. epidermidis* over a 7-day period (n = 8 per group). For this survival analysis, the terminal endpoint was defined as complete barrier failure, evidenced by the translocation of bacteria into the basal culture medium. This breakthrough contamination was monitored daily and confirmed by either macroscopic medium turbidity or the growth of viable colonies from basal medium aliquots (5 × 10 μL per model). (D) Representative macroscopic images and corresponding H&E histological cross-sections of the HEEs treated with PBS (control), *S. epidermidis*, *P. aeruginosa*, and MRSA at day 2, 4, and 7. Scale bars, 20 μm. Red arrowheads highlight localized bacterial clusters.

In contrast, the opportunistic pathogens proliferated without restriction. The bacterial burdens of both *P. aeruginosa* and MRSA increased from approximately 10^6^ to 10^7^ CFU/model at day 2 to more than 10^9^ CFU/model by day 7 (Figure 2B). Relative to the commensal baseline, this difference was statistically significant as early as day 2 (*p* < 0.05 for *P. aeruginosa*, *p* < 0.0001 for MRSA) and increased further by day 4 and day 7 (*p* < 0.0001 for both strains). The increase in bacterial burden was accompanied by rapid tissue collapse. The skin survival rate in the *P. aeruginosa*-infected group began to decline by day 6 and fell below 50% by day 7, whereas the MRSA-infected models exhibited approximately 85% survival at day 7 (Figure 2C). Because model failure was defined as translocation of bacteria into the basal medium, these survival values also define the proportion of models in which apical–basolateral separation was still intact at the point of sampling; the consequences of this for the day-7 secretome analysis are considered in the Discussion. Macroscopic observations and histological evaluations of H&E-stained sections closely mirrored this temporal trajectory (Figure 2D). While the structural architecture remained relatively intact at day 2, the models underwent progressive epidermal necrosis, resulting in the widespread detachment of the stratum corneum and underlying viable layers by day 7.

### 2.3. 3D Spatial and Microstructural Characterization of Bacterial Invasion

To quantify the penetrative capacity of pathogens into tissue depths, we performed *in situ* whole-mount confocal imaging on HEEs infected with fluorescently tagged strains and measured the apparent invasion depth from the outermost keratinocyte layer (see Methods in detail). On day 2, all strains were largely confined to the epidermal surface (apparent invasion depth ∼ 0 - 1 μm). By day 7, the commensal *S. epidermidis* remained restricted to the superficial layer (<1 μm) whereas the opportunistic pathogens penetrated the tissue: *P. aeruginosa* to an apparent depth of approximately 13 μm (*p* < 0.001), and MRSA reached approximately 14 μm (*p* < 0.001) (Figure S1). High-resolution 3D volume renderings visually corroborated these measurements, showing that the pathogens breached the stratum corneum and assembled dense three-dimensional microcolonies within the viable cellular layers (**Figure 3**).

**Figure 3.**
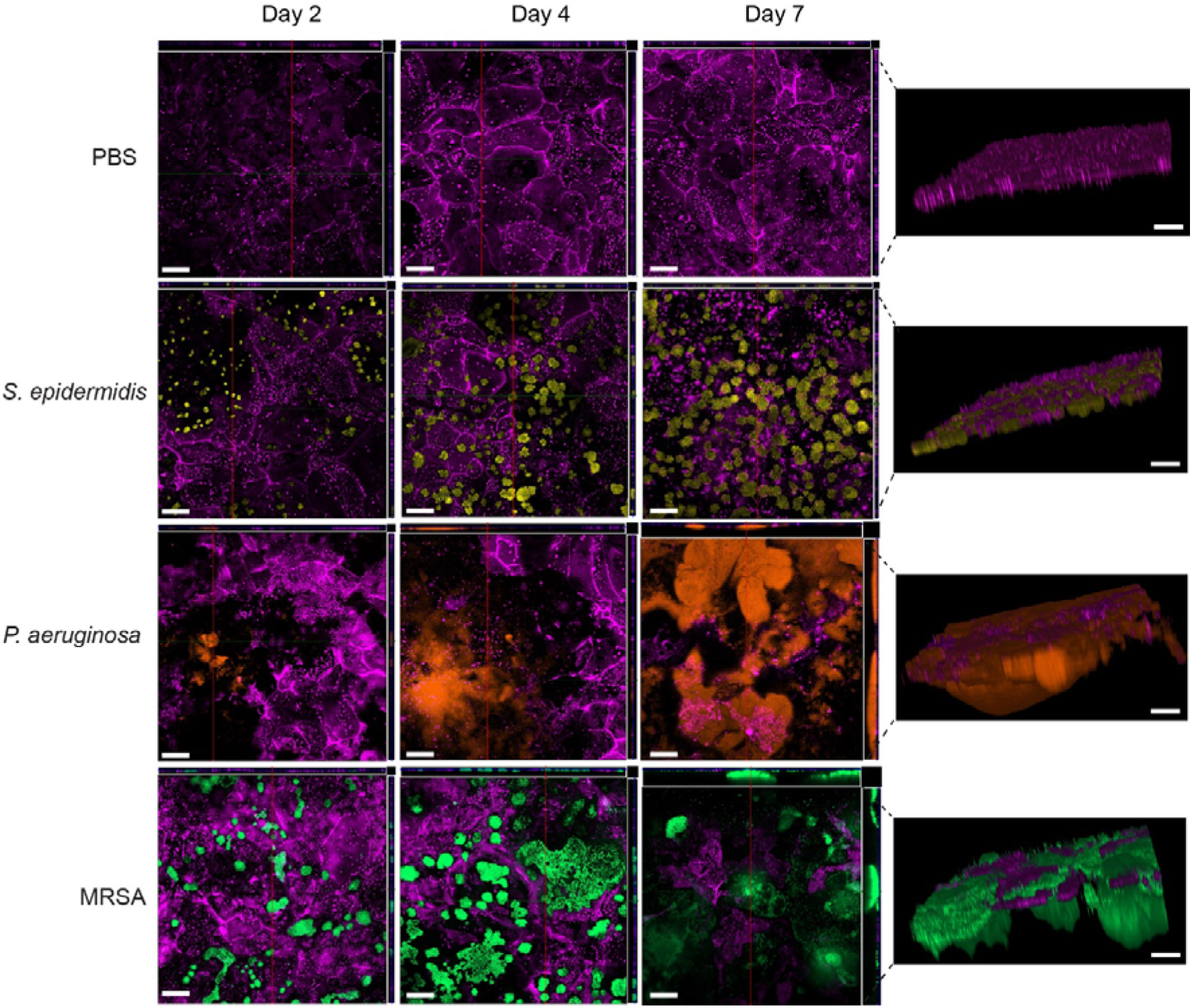
3D spatial mapping and quantification of progressive bacterial epidermal invasion. Representative whole-mount confocal microscopy images illustrating the spatial distribution of fluorescently tagged bacteria within the epidermal equivalents at day 2, 4, and 7. Left panels show the maximum intensity projections (XY plane), while right panels display the corresponding 3D volume renderings. Scale bars, 50 μm. *Note: Host keratinocyte membranes are shown in magenta; note that S. epidermidis and P. aeruginosa both carry mCherry and MRSA carry GFP. For optimal visualization of the spatial invasion depth, the Z-axis in the 3D volume renderings was digitally scaled by a factor of 5 (from the original 2* μ*m physical Z-step to 10* μ*m per slice) relative to the X and Y axes.*

Scanning electron microscopy (SEM) provided further morphological alterations associated with this invasive phenotype. Surface topography revealed a clear transition from scattered, individual bacteria at day 2 to expansive, dense biofilm-like networks by day 7, particularly pronounced within the *P. aeruginosa* and MRSA groups (Figure S2). Concurrently, cross-sectional SEM imaging explicitly showed the collapse of the physical barrier (**Figure 4**). The initially cohesive, multi-layered stratum corneum observed in the control and commensal groups was delaminated and degraded in the pathogen-exposed groups, most conspicuously in *P. aeruginosa*-infected models, in which the viable layers were converted into a highly porous, honeycomb-like structure by day 7. This structural dissolution is consistent with invasion observed in the whole-mount staining images.

**Figure 4.**
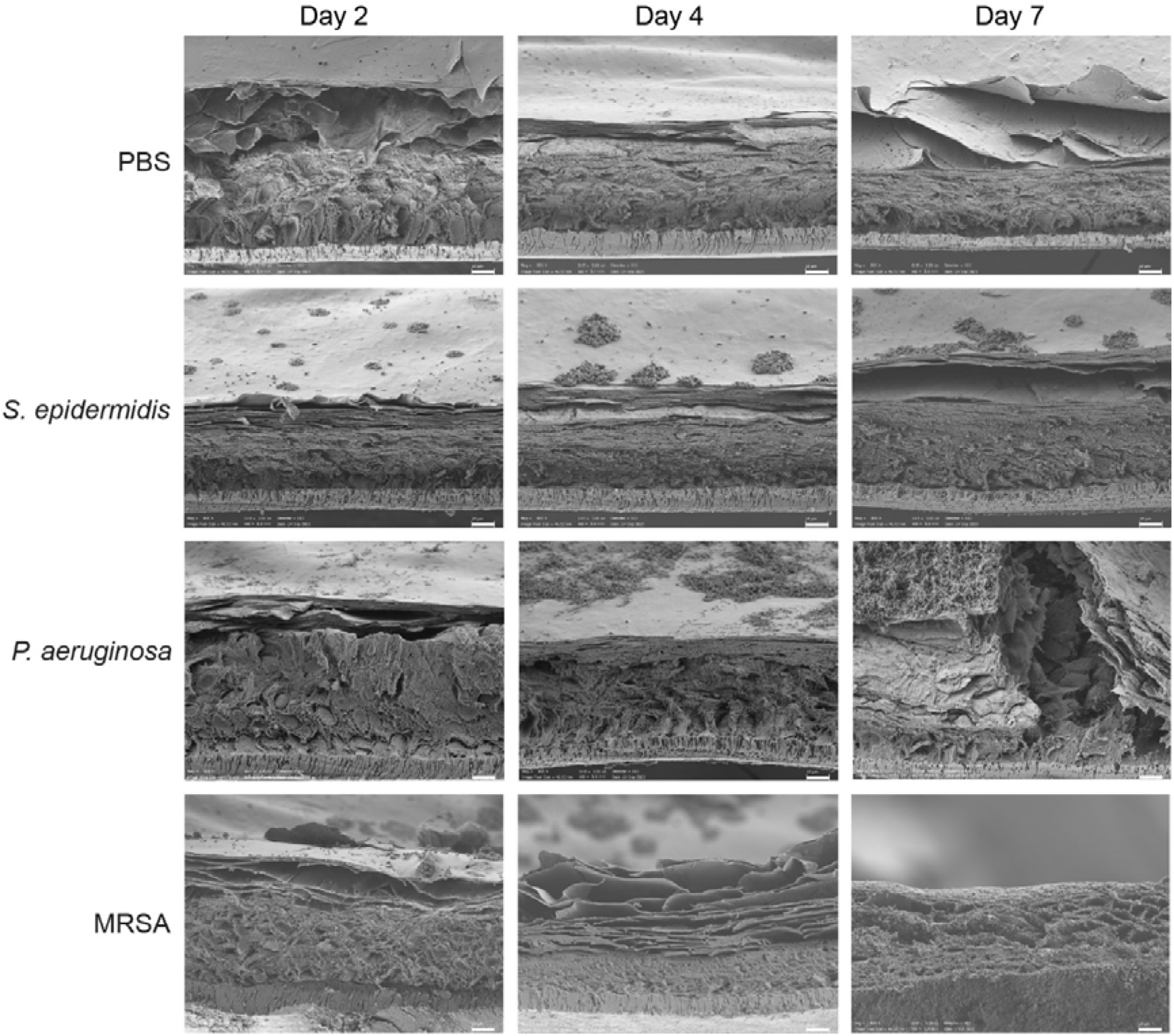
Cross-sectional morphological analysis of pathogen-induced epidermal barrier disruption. Representative cross-sectional scanning electron microscopy (SEM) images of HEEs treated with PBS, *S. epidermidis*, *P. aeruginosa*, and MRSA at day 2, 4, and 7. Scale bars, 10 μm.

### 2.4. Longitudinal proteomic profiling identifies IL-1α as a damage-associated readout of epidermal infection

To characterize host epidermal responses to different bacterial species and identify biomarkers associated with the transition from colonization to tissue disruption, we performed longitudinal proteomic profiling using the Olink platform on basolateral conditioned media collected from four experimental groups at day 2, 4, and 7. Principal component analysis (PCA) showed no distinct proteomic separation in the commensal *S. epidermidis* group over the 7-day period, which clustered closely with the PBS controls (**Figure 5A**). In contrast, both the *P. aeruginosa* and MRSA groups diverged markedly on day 7 compared to day 2 and day 4, coinciding with the onset of structural barrier damage. Longitudinal fold-change trajectories relative to the day 2 baseline further highlighted distinct regulatory patterns between the two pathogens (Figure 5B). Specifically, *P. aeruginosa* infection induced a bidirectional pattern by day 7, characterized by both strongly upregulated (e.g., IL-18R1) and downregulated proteins (e.g., IL-10RB, CCL19, and TRAIL; Figure 5B, Figure S3A). Conversely, MRSA infection predominantly drove a broad upward trend across the profiled markers by day 7 (e.g., selective induction of 4E-BP1, IL-12B, IL-17C, and TNF; Figure 5B, Figure S3B).

**Figure 5.**
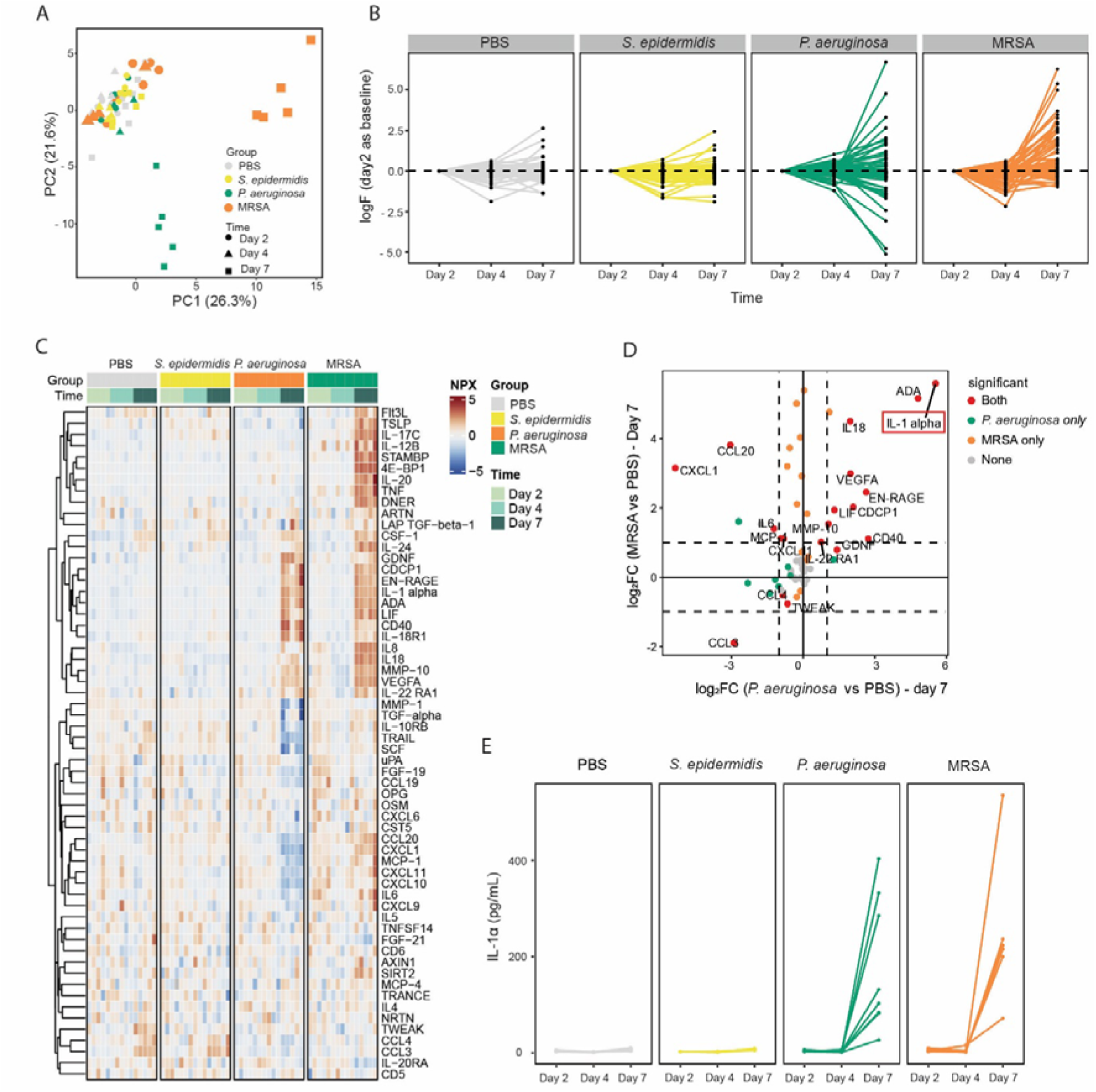
Longitudinal proteomic profiling of basolateral conditioned media during epidermal infection. (A) Principal component analysis (PCA) of profiles of 60 inflammation-related proteins across four groups (PBS, *S. epidermidis*, *P. aeruginosa*, and MRSA) at days 2, 4, and 7 (n = 5 per group). (B) Temporal trajectories of protein abundance within each group, shown as mean log2 fold change (log_2_FC) relative to the day 2 baseline. Each line represents a single protein. (C) Heatmap showing hierarchical clustering of Normalized Protein eXpression (NPX) values for 60 inflammation-related proteins across groups and time points. (D) Scatter plot comparing log_2_FC at day 7 (relative to PBS) between *P. aeruginosa* and MRSA groups. Dashed lines indicate a |log_2_FC| threshold of 1. Colors denote statistical significance (adjusted *p* < 0.05). IL-1α is highlighted in the red box. (E) Absolute IL-1α concentrations (pg/mL) in conditioned media at days 2, 4, and 7 quantified by ELISA (n = 5-8 per group). Lines connect repeated measurements from individual models.

Consistent with these trajectories, hierarchical clustering illustrated both strain-specific inflammatory signatures and shared response modules across the bacterial groups (Figure 5C). While investigating strain-specific responses is valuable, we aimed to identify a broad-spectrum biomarker that could signal infection regardless of the causative species. Since *P. aeruginosa* and MRSA serve as representative models for Gram-negative and Gram-positive skin pathogens^[22]^, markers commonly upregulated by both strains may reflect shared host responses to opportunistic bacterial infection. Therefore, we analyzed the proteins significantly altered at day 7 in either or both infection groups relative to PBS controls (Figure 5D). Among the core subsets of co-upregulated proteins, including ADA, IL-18, EN-RAGE, and VEGFA (Figure S3C), IL-1α emerged as the most prominent shared indicator (Figure 5D).

To verify these proteomic findings and evaluate the suitability of IL-1α for bioelectronic sensing, we quantified absolute IL-1α concentrations in the conditioned media by ELISA (Figure 5E). Confirming the screening data, IL-1α levels remained near baseline from day 2 to day 4 across all groups, as well as at day 7 in the PBS and *S. epidermidis* controls, but rose significantly between day 4 and day 7 in both the *P. aeruginosa* and MRSA groups. This low-background elevation indicates that, within this model, IL-1α provides a readout of the shift from superficial colonization to active epidermal damage.

### 2.5. Development of a flexible electrochemical immunosensor for IL-1α screening

The infection response identified in the 3D epidermal models, together with the Olink-based inflammatory profiling, provided the basis for selecting IL-1α as the molecular target for electrochemical sensing. Its distinct secretion profile and pronounced increase during infection supported its evaluation as a candidate biomarker for distinguishing different infection states. To explore the potential for rapid screening, we developed a flexible electrochemical immunosensor based on Au electrodes fabricated on 75-µm-thick polyimide (PI) substrates by radiofrequency (RF) magnetron sputtering, followed by laser patterning of the three-electrode geometry. SEM confirmed the formation of a continuous and homogeneous Au coating (Figure S4). Initial electrochemical characterization of the bare electrodes using the Fe(CN_6_)^3-/4-^redox probe revealed good inter-device reproducibility with a relative standard deviation (RSD) of 4% (Figure S5A-B), stable scan-to-scan responses, and predominantly diffusion-controlled electrochemical behavior (Figure S5C). Furthermore, the electrochemical response increased proportionally with redox probe concentration (Figure S5D), confirming the suitability of the fabricated electrodes for subsequent biofunctionalization.

Next, the Au working electrode was functionalized using a mixed 11-mercaptoundecanoic acid/6-mercapto-1-hexanol (MUA/MCH) self-assembled monolayer, followed by EDC/NHS activation, immobilization of anti-IL-1α antibodies and bovine serum albumin (BSA) blocking (**Figure 6A-B**). Stepwise electrochemical impedance spectroscopy (EIS) measurements showed progressive increases in charge-transfer resistance following surface modification and subsequent IL-1α binding, consistent with changes in electron transfer at the electrode interface (Figure 6C-D). The immunosensor exhibited a concentration-dependent response to IL-1α over the investigated concentration range of 25–200 pg/mL (Figure 6E). Fitting the mean normalized response against log10[IL-1α] gave a slope of 0.906 ± 0.009 and a coefficient of determination (R² = 0.9998) (Figure 6E). The estimated limit of detection (LOD) and limit of quantification (LOQ) were 2.93 and 26.09 pg/mL, respectively. Inter-electrode variability evaluated using three independently prepared electrodes, yielded RSD values ranging from 9.2% to 34% across the tested concentrations. Furthermore, the sensor demonstrated selectivity toward IL-1α in the presence of other inflammatory cytokines (Figure 6F).

**Figure 6.**
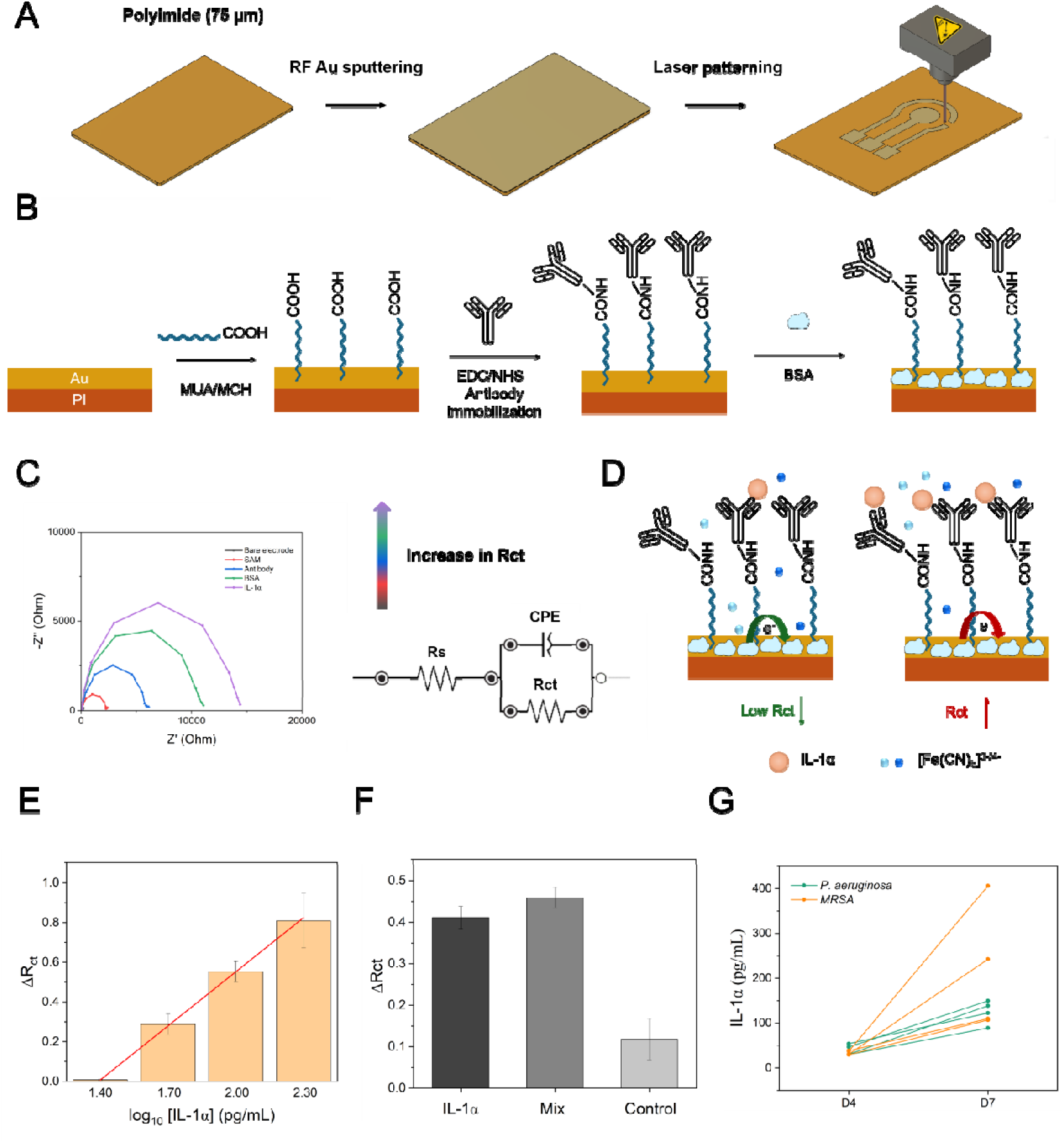
Fabrication, biofunctionalization, sensing mechanism, and performance characterization of the Au-based electrochemical sensor for IL-1α detection. **(A)** Fabrication of the flexible electrode by RF sputtering of Au onto a 75-μm-thick polyimide substrate, followed by laser patterning to define the electrode architecture. **(B)** Surface functionalization of the Au electrode through MUA/MCH self-assembled monolayer formation, followed by EDC/NHS-mediated immobilization of antibodies and subsequent blocking with BSA. **(C)** Representative electrochemical impedance spectra and equivalent circuit used to characterize the electrode interface at different modification stages. The circuit comprises the solution resistance (R_s_), constant phase element (CPE), and charge-transfer resistance (R_ct_); increasing R_ct_ indicates progressive interfacial modification. **(D)** Proposed sensing mechanism based on target-induced changes in interfacial electron transfer. Binding of the target species alters the accessibility of the redox probe Fe(CN)□³□/□□to the electrode surface, resulting in a measurable change in. (E) Concentration-response of the immunosensor, showing normalized EIS charge-transfer resistance (ΔRct) for increasing IL-1α concentrations (25, 50, 100, and 200 pg/mL) obtained using three independent electrodes (mean ± SD). The line shows the linear regression of the mean response against log10[IL-1α]; the response at 25 pg/mL was not resolved from the pre-exposure baseline. (F) Selectivity assessment of the developed sensor for IL-1α in the presence of a mixture of potential interfering cytokines, IL-1β, IL-6, and IL-8, compared with the control without IL-1α. Measurements were performed using three independent electrodes for each condition (n = 3). (G) Proof-of-concept evaluation using conditioned media from infected HEE models. Green, *P. aeruginosa* samples collected at days 4 and 7; orange, MRSA samples collected on days 4 and 7. Sensor-derived IL-1α concentration equivalents increased from day 4 to day 7 for both infection models. Values above 200 pg/mL lie outside the calibrated range and are extrapolated; conditioned media from the PBS and *S. epidermidis* groups were not measured.

As proof of concept, the flexible immunosensor was evaluated using conditioned media collected from previously infected HEE models exposed to *P. aeruginosa* and MRSA. Previous characterization of these infection models showed a marked temporal increase in IL-1α secretion during infection, providing a biological context to assess whether the flexible immunosensor could resolve changes in infection state over time. Conditioned-media samples collected on days 4 and 7 of infection were analyzed using the flexible immunosensor. The calibration relationship between normalized electrochemical response and log_10_[IL-1α] was used to express the measured responses as sensor-derived IL-1α concentration equivalents. For *P. aeruginosa*-infected HEE models, the derived values increased from 29.8–54.2 pg/mL at day 4 to 89.2–149.7 pg/mL at day 7. For MRSA-infected models, the corresponding values increased from 30.7–38.6 pg/mL to 107.1–406.1 pg/mL (Figure 6G). Thus, both infection models exhibited higher sensor responses on day 7 than on day 4, with a particularly pronounced increase in the MRSA-infected samples.

## 3. DISCUSSION

Here, we established a standardized 3D human epidermal infection platform to identify host signals of tissue-destructive infection, and selected one of them, IL-1α, as the target of a flexible Au/PI immunosensor. In our longitudinal model, bacterial proliferation outpaced the host inflammatory response. Although *P. aeruginosa* and MRSA were already abundant at day 2, the proteomes at days 2 and 4 remained indistinguishable from PBS and *S. epidermidis* controls, diverging only at day 7. The stratum corneum is a plausible explanation for this delay. In stratified epidermis, surface bacteria are separated from viable keratinocytes, thereby limiting their direct exposure to superficial microorganisms.^[23,24]^. Our longitudinal characterization data supports this timeline: while all strains were confined to the surface at day 2, pathogenic strains had breached the stratum corneum by day 7, forming dense microcolonies and biofilm-like networks within viable layers alongside a delaminated, porous barrier, coinciding with the observed proteomic shift. As a result, surface bacterial burden alone cannot reliably indicate when an epidermal tissue has crossed this structural threshold from superficial colonization to active invasion, highlighting the need for host-derived biomarkers that directly reflect barrier breakdown.

*P. aeruginosa* and MRSA triggered different proteomic trajectories at day 7, which complicates biomarker selection. MRSA raised a broad set of inflammatory mediators, including TNF, IL-8 and IL-12B, whereas *P. aeruginosa* gave a mixed picture in which some proteins rose and others, such as IL-10RB, CCL19 and TRAIL, fell. This difference aligns with the distinct pathogenic strategies of the two species. *S. aureus* is sensed through innate pathways, including TLR2 and the inflammasome, that drive new cytokine transcription^[25]^. By contrast, *P. aeruginosa* secretes exotoxin A, which blocks host protein synthesis^[26]^, and proteases such as elastase (LasB) and alkaline protease (AprA) that cleave cytokines and chemokines to evade immune clearance^[27]^. These pathogen-specific mechanisms may contribute to differences in the abundance of inflammatory proteins detected in the conditioned media. For biosensor development, this reinforces the importance of selecting biomarkers whose levels are less dependent on pathogen-specific host responses or susceptible to bacterial degradation, potentially enabling more consistent detection across different bacterial infections.

A marker less exposed to this variation would be one released by injury and not produced in response to it. IL-1α fits that description and was the most consistent shared increase in both pathogen groups. Unlike inducible cytokines that require active de novo synthesis by viable cells, IL-1α is constitutively synthesized and stored as a biologically active precursor within the cytoplasm of healthy keratinocytes^[28]^. Upon membrane disruption and cellular necrosis, this preformed pool is passively released into the extracellular space, a mechanism well established in reconstructed human epidermis irritation assays^[29]^. Because both *P. aeruginosa* and MRSA cause epidermal necrosis and barrier disintegration, the passive release of preformed IL-1α bypasses pathogen-specific immune modulation, providing a clear, low-background signal of active tissue disruption. We emphasize that IL-1α release from reconstructed human epidermis is itself an established irritation endpoint; what the present screen adds is that, among 60 inflammation-related proteins measured in parallel, IL-1α was the most consistent shared responder across a Gram-negative and a Gram-positive pathogen, and that the mechanistic basis for that consistency is precisely the property that recommends it as a sensing target in a protease-rich matrix.

Having identified IL-1α as a shared marker of infection-associated tissue damage, we next developed a flexible electrode platform to translate its recognition into an electrochemical signal. The platform combines thin-film metallization of a polyimide (PI) substrate with digitally defined laser patterning. This separation of metal deposition and electrode patterning allows electrode dimensions and layouts to be modified digitally without requiring new lithographic masks, conductive ink formulations or printing templates. The approach therefore facilitates rapid prototyping while combining the electrical properties of Au with the flexibility of PI^[30,31]^, a material widely used in flexible electronics^[32,33]^. Although wearable performance and integration were not evaluated in this study, the fabrication strategy provides a basis for future skin-interfaced or portable electrochemical platforms.

The as-fabricated Au electrodes exhibited an inter-device RSD of approximately 4% within a single fabrication batch. However, variations across sputtering batches and substrate positions may reflect differences in deposited film thickness, surface morphology and microstructure ^[34,35]^, potentially affecting their initial electrochemical properties and subsequent functionalization. Responses were therefore normalized to each electrode’s pre-assay baseline to reduce the influence of differences in initial electrochemical behavior. Although this approach facilitates comparisons between electrodes, it cannot compensate for variability introduced during biofunctionalization. Accordingly, the complete immunosensor exhibited higher inter-electrode RSD values, ranging from 9.2% to 34% across the investigated concentrations (n = 3). These findings suggest that, beyond improving deposition consistency, further standardization of the biofunctionalization procedure will be necessary to improve the reproducibility of the final sensing interface.

The observed increases in charge-transfer resistance following successive functionalization steps, including IL-1α exposure, were consistent with formation of the intended immunosensing interface. The concentration-dependent response and initial selectivity assessment against IL-1β, IL-8 and IL-6 further supported target recognition, although more extensive validation is required. The estimated limit of quantification (LOQ) was 26.09 pg/mL, slightly above the lowest investigated concentration of 25 pg/mL. Quantification is therefore more appropriately considered feasible at or above the estimated LOQ, pending further validation. The sensor was subsequently evaluated using conditioned media from the infected HEE models. In agreement with the ELISA measurements, which showed a pronounced increase in IL-1α between days 4 and 7, the electrochemical responses were higher for day 7 than day 4 samples from both pathogen groups. This difference was particularly pronounced for MRSA but was also evident for *P. aeruginosa.* These findings demonstrate the ability of the sensor to distinguish conditioned media associated with different stages of infection-related tissue responses within the pathogen-exposed groups. However, the calibration range was limited to 25–200 pg/mL, and responses approached saturation above this range. Consequently, values exceeding 200 pg/mL, including day 7 MRSA measurements, should be regarded as extrapolations rather than validated quantitative estimates. The electrochemical results therefore demonstrate state discrimination rather than quantitative agreement with ELISA or replacement of established cytokine measurements.

However, several limitations should be considered when interpreting these findings and planning further development. The HEE model relies on a single immortalized keratinocyte line and lacks immune cells, fibroblasts and vasculature. It therefore primarily reflects keratinocyte-intrinsic responses and does not capture immune-cell recruitment. Furthermore, IL-1α increased alongside overt structural damage rather than clearly before it. More closely spaced sampling between days 4 and 7 will be required to establish the temporal relationship between IL-1α release and barrier failure and to determine whether the biomarker can provide an earlier indication of tissue disruption. The sensor’s investigated calibration range (25–200 pg/mL) is also narrower than the IL-1α concentrations reported in chronic wound fluid (340–11,000 pg/mL) in leg ulcers without evident clinical signs of infection^[36]^. Concentrations in infected human wounds remain to be established, although an ex vivo human skin model showed increased IL-1α following *P. aeruginosa* infection^[37]^. Expanding the sensor’s dynamic range, evaluating its performance in complex wound matrices and establishing suitable sample dilution protocols will therefore be important. Finally, improved control of electrode deposition and biofunctionalization, together with testing of larger numbers of independently fabricated devices and biological samples, will be required to address inter-electrode variability. The present evaluation also relied on collected conditioned media at defined endpoints, rather than continuous monitoring or direct measurements at the tissue–sensor interface.

Taken together, this study establishes a framework in which tissue-level infection biology guides the development of flexible electrochemical sensing. The HEE model identified IL-1α as a shared host-derived signal associated with the transition from superficial colonization to tissue invasion and damage. The flexible Au/PI immunosensor demonstrated the feasibility of resolving infection-stage-dependent changes in those media. Although further work is needed to improve reproducibility, to quantify and subtract the matrix-associated background, to expand the dynamic range and to establish performance in clinically relevant samples, this approach provides a basis for developing sensing platforms informed by, and tested against, the biological processes they are intended to monitor.

## 4. MATERIALS AND METHODS

### 4.1. Materials and Reagents

All chemical reagents and materials were used as received without further purification. Polyimide (PI) films (75 μm thickness) and double-sided adhesive tape were purchased from RS Components. The 3M 9969 transfer adhesive medical tape was provided by 3M. Isopropyl alcohol (IPA), 6-mercapto-1-hexanol (MCH), 11-mercaptoundecanoic acid (MUA), absolute ethanol, 1-ethyl-3-(3-dimethylaminopropyl)carbodiimide hydrochloride (EDC·HCl, N-hydroxysuccinimide (NHS), 2-(N-morpholino)ethanesulfonic acid (MES), bovine serum albumin (BSA), Ag/AgCl conductive ink (60/40 formulation), potassium ferrocyanide trihydrate, potassium ferricyanide, and phosphate-buffered saline (PBS) were all purchased from Sigma-Aldrich. Recombinant human cytokines were sourced as follows: IL-1α (Fisher Scientific), IL-1β (ImmunoTools), IL-6 (PromoCell), and IL-8 (R&D Systems/Bio-Techne). Commercial human ELISA kits were purchased from R&D Systems. The Ag/AgCl reference electrode (CHI111) was obtained from CH Instruments. CellBrite™ Red Cytoplasmic Membrane Dye was purchased from Biotium. Milli-Q water (resistivity of 18.2 MΩ·cm) was used for all aqueous solution preparations.

### 4.2. Bacteria Growth and Preparation

Fluorescently tagged bacterial strains, including GFP-expressing *Staphylococcus aureus* (USA300), mCherry-expressing *Pseudomonas aeruginosa* (PAO1), and mCherry-expressing *Staphylococcus epidermidis*, were preserved as glycerol stocks at −80 °C. Two days prior to HEE inoculation, bacterial stocks were streaked onto Tryptic Soy Agar (TSA) plates supplemented with appropriate antibiotics to maintain plasmid selection and incubated overnight at 37 °C. A single colony of each strain was then inoculated into Tryptic Soy Broth (TSB) containing respective antibiotics and cultured overnight at 37 °C with continuous shaking at 200 rpm. On the day of infection, overnight cultures were spectrophotometrically adjusted to standardize the initial bacterial concentration. The OD600 was adjusted to 0.1 for *P. aeruginosa* and 0.3 for *S. aureus* and *S. epidermidis*, corresponding to a uniform density of approximately 1 × 10^8^ CFU/mL across all strains. The suspensions were serially diluted in sterile PBS to a working concentration of 1 × 10^6^ CFU/mL for topical inoculation.

### 4.3. Keratinocyte Culture and Generation of Epidermal Equivalents

N/TERT-2G keratinocytes were maintained in EpiLife medium^[38]^. Human epidermal equivalents (HEEs) were generated on type I collagen-coated polycarbonate inserts by seeding 2 × 10^5^ N/TERT-2G cells per insert. Cultures were raised to the air–liquid interface after 3 days and maintained in a medium supplemented with 1.5 mM CaCl□ and 50 μg·mL^-1^ vitamin C. Mature HEEs (12 days post-airlift) were used for infection modeling.

### 4.4. Bacteria Inoculation and Survival Monitoring on HEEs

Topical bacterial colonization was performed using a standardized glass cloning cylinder methodology to restrict the infection area. Mature HEEs (with or without prior 0.1% SDS treatment for 30 min, depending on acute/chronic experimental designs) were placed on a 37 °C heating plate within a biosafety cabinet. Glass cloning cylinders (outer diameter 6 mm, inner diameter 4 mm; Bioptechs, USA), sterilized with ethanol, were positioned centrally on the HEE apical surface (stratum corneum). Subsequently, 25 μL of the diluted bacterial suspension (yielding an initial inoculum of ∼2.5 × 10^4^ CFU/model) or sterile PBS (mock control) was pipetted into the cylinder. Cultures were left unlidded on the heated plate for 4–5 h to allow complete evaporation of the suspension. The cylinders were then removed, basal media were replenished if necessary, and models were incubated (37 °C, 5% CO_2_). *Model Survival Assessment:* For Kaplan-Meier survival analysis, the terminal endpoint of the HEE model was strictly defined as complete barrier failure, evidenced by the translocation of pathogens into the basal culture medium. This breakthrough contamination was monitored daily and confirmed by either macroscopic medium turbidity or the growth of viable colonies following the agar plating of basal medium aliquots (10 μL, 5 replicates per model).

### 4.5. TEER Measurement

Transepithelial electrical resistance (TEER) was measured using an EVOM3 system equipped with chopstick electrodes. A standardized volume of sterile PBS was gently applied to the apical surface of the HEEs to establish electrical continuity. Values were corrected for cell-free blank inserts, normalized to the effective membrane surface area, and expressed as Ω·cm^2^.

### 4.6. CFU Counting

To quantify the bacterial burden, HEEs were harvested at specified time points post-inoculation. Epidermal tissues were excised, transferred into sterile PBS, and mechanically homogenized. Tissue homogenates were serially diluted and plated onto LB agar. Following overnight incubation at 37 °C, colonies were counted, and the total bacterial load was expressed as CFU per epidermis model.

### 4.7. Histological Analysis (H&E)

Harvested HEEs were fixed in 4% paraformaldehyde (PFA) for ≥24 h at room temperature, dehydrated through a graded ethanol series, cleared in xylene, and embedded in paraffin wax. Tissue blocks were sectioned transversely (10 μm thickness). Sections were deparaffinized, rehydrated, and stained with hematoxylin and eosin (H&E) using standard protocols to evaluate morphological alterations and barrier integrity under a bright-field microscope.

### 4.8. Scanning Electron Microscopy (SEM)

For morphological analysis, infected HEEs were immersion-fixed in 2.5% glutaraldehyde prepared in 0.1 M phosphate buffer (pH 7.4). Samples were washed with Milli-Q water, stepwise dehydrated in ethanol, and critical-point dried using carbon dioxide (Leica EM CPD 030). Dried samples were mounted on pin stubs with conductive carbon tabs and sputter-coated with platinum (Quorum Q150T ES). SEM was performed using a Zeiss Ultra 55 field-emission scanning electron microscope operated at an accelerating voltage of 3 kV and equipped with an SE2 detector. Images were acquired to examine surface biofilm morphology and the microstructural organization of cross-sectional tissue barrier architectures.

Gold electrodes were also examined by SEM using the same microscope settings. As the electrodes were already gold-coated and electrically conductive, no additional sputter coating was applied.

### 4.9. In Situ Live Whole-Mount Confocal Imaging and 3D Invasion Quantification

To visualize the 3D spatial distribution and quantify the penetration depth of live bacteria in their native state, HEEs infected with fluorescent-tagged strains were subjected to whole-mount imaging. At indicated time points, infected HEEs were harvested and washed with PBS. Epidermal cytoplasmic membranes were stained with CellBrite™ Red Cytoplasmic Membrane Dye (Biotium) according to the manufacturer’s protocol. Notably, chemical fixation (e.g., PFA) and detergent permeabilization were deliberately omitted to prevent the quenching of bacterial fluorescent proteins and to strictly preserve the native 3D architecture of the host-pathogen interface. Intact models were mounted onto 25 mm confocal glass-bottom dishes and immediately imaged using an inverted confocal laser scanning microscope (CLSM). High-resolution Z-stack images were acquired from the stratum corneum down to the basal layers.

#### Quantification of Invasion Depth

Bacterial epidermal invasion depth was quantified using ZEN/ImageJ software. The depth was calculated as the Z-axis orthogonal distance between the uppermost layer of the keratinocytes (Z=0 surface) and the deepest boundary of the bacterial fluorescent signal, based on the 2 μm Z-step interval. For each experimental group, 5 independent HEEs were analyzed. For each individual HEE model, a representative field of view (FOV) was imaged, and the invasion depths of 3 distinct bacterial microcolonies were measured and averaged to yield a single mean value per HEE prior to inter-group statistical comparisons. Since fluorescence attenuates with imaging depth in unfixed whole mounts and no independent marker of the stratum corneum/viable-layer boundary was used, these values are likely to underestimate true penetration.

#### 3D Volume Rendering

For the generation of 3D volume renderings, the Z-axis aspect ratio was digitally adjusted (scaled by 5×, assigning 10 μm per actual 2 μm slice) to overcome the visual flattening effect inherent to large XY-field scans of thin tissue sections, thereby optimizing the visualization of the spatial relationship between invading pathogens and host tissue.

### 4.10. Targeted proteomic profiling and statistical analysis

Longitudinal proteomic profiling of basolateral conditioned media (1 μL per sample) was performed using the Olink Target 96 Inflammation panel on the Olink Signature Q100 platform (Olink Proteomics, Uppsala, Sweden) according to the manufacturer’s instructions. The dataset comprised 60 samples across four groups (PBS, *S. epidermidis*, *P. aeruginosa*, and MRSA; n = 5 biological replicates per group) collected at days 2, 4, and 7. Protein abundance was quantified as Normalized Protein eXpression (NPX, log2 scale) using NPX Signature software (version 2.2.1). Following quality control, assays with values below the limit of detection (LOD) in ≥ 95% of samples were excluded, leaving 60 out of 92 proteins for downstream analysis.

All statistical analyses were conducted in R. Principal component analysis (PCA) was performed on centered and unit-variance-scaled NPX values using FactoMineR (v2.12). Temporal trajectories were visualized using mean log_2_ fold changes (log_2_ FC) relative to the day 2 baseline. To evaluate group-specific temporal changes while accounting for repeated measurements, linear mixed-effects models (NPX ∼ time * group + (1 | Individual)) were fitted via restricted maximum likelihood (REML) using lmerTest (v3.2.1), and the group-by-time interaction was assessed by Type III ANOVA with Satterthwaite’s approximation. Model_ID identifies the individual epidermal equivalent from which basolateral medium was sampled repeatedly at days 2, 4 and 7; CFU counting, histology and electron microscopy required destructive harvest and were therefore performed on separate models. Differential protein abundance at day 7 relative to the PBS control was analyzed using limma (v3.64.1; lmFit and eBayes functions). Across all analyses, p-values were adjusted for multiple testing using the Benjamini-Hochberg (BH) method, with an adjusted p < 0.05 considered statistically significant.

### 4.11. ELISAs

The secretion of IL-1α was quantified from the basal culture medium. Supernatants were collected, centrifuged to remove cellular debris, and measured using commercial ELISA kits (R&D Systems). The optical density was measured at 450 nm, and concentrations were calculated against standard curves.

### 4.12. Fabrication of Flexible Au/PI Electrodes

Flexible electrodes were fabricated on 75 μm-thick PI substrates. The electrode geometries were designed digitally using Autodesk Fusion CAD software. A titanium/tungsten (Ti/W) adhesion layer ^[39]^ followed by approximately 100 nm of Au was deposited by RF magnetron sputtering. The electrode geometries were subsequently defined by direct laser patterning using a desktop laser instrument. Before functionalization, the electrodes were rinsed with Milli-Q water and IPA, gently wiped with KIMWIPES, and dried under ambient conditions. Double-sided adhesive tape was also applied to the back of the electrodes to improve handling during subsequent experimental procedures. The surfaces were then treated with air plasma for 30 s at 100 W. A laser-patterned 3M 9969 transfer adhesive mask was used to define the exposed areas of the working and reference electrodes. Then, the Ag/AgCl reference electrode was defined using Ag/AgCl conductive ink. The ink was applied to the designated reference-electrode area and cured at 80 °C for 10 min in an oven. Its electrochemical performance was then evaluated by comparison with a standard external reference electrode to verify its behaviour.

### 4.13. Sensor Surface Functionalization

The Au working electrode was modified with a mixed self-assembled monolayer (SAM) of MUA and MCH. MUA (0.5 mM) and MCH (1.0 mM) were prepared in absolute ethanol at a 1:2 molar ratio. Electrodes were incubated in the SAM solution for 16 h at room temperature in the dark and subsequently rinsed with ethanol and Milli-Q water. Carboxyl groups from MUA were activated using 0.2 M EDC·HCl and 0.05 M NHS in 0.1 M MES buffer (pH 5.0) for 30 min at room temperature^[40]^. After rinsing, anti-IL-1α antibodies (50 μg/mL) prepared in 10 mM PBS (pH 7.4) were immobilized for 1 h at room temperature. Following antibody immobilization, the electrodes were blocked with 1% BSA in 10 mM PBS (pH 7.4) for 60 min to reduce nonspecific adsorption. The electrodes were subsequently rinsed with PBS and stored at 4 °C in PBS until use.

### 4.14. Electrochemical Measurements

The flexible electrodes were interfaced with the potentiostat using a Gii-Sens SPE Adapter holder (PalmSens). Measurements were performed using a μStat-i 400s potentiostat (Metrohm). EIS spectra were recorded from 10 kHz to 0.05 Hz using a 10 mV AC perturbation. The open-circuit potential was monitored for 60 s, followed by a 30 s equilibration period. Measurements were performed using a redox-probe solution containing 5 mM [Fe(CN)_6_]³ and 5 mM [Fe(CN)_6_] in 10 mM PBS (pH 7.4). The EIS spectra were fitted using an equivalent electrical circuit consisting of the solution resistance (Rs) in series with a parallel combination of a constant-phase element (CPE) and charge-transfer resistance (Rct), i.e., Rs – (CPE || Rct). High-frequency data points exhibiting inductive behavior (-Z’’<0) were excluded from the fitting procedure to minimize the influence of external wiring contributions and avoid distortion of the fitted parameters. For IL-1α measurements, conditioned-media samples or IL-1α solutions were applied to the functionalized working electrode for 10 min, followed by rinsing with PBS before EIS measurement. To account for differences in the baseline electrochemical properties of individual electrodes, the response was expressed as the normalized charge-transfer resistance response: ΔR_ct_ = (R_ct_ - R_ct,0_)/R_ct,0_, where R_ct,0_ is the baseline charge-transfer resistance measured immediately before sample exposure and R_ct_ is the charge-transfer resistance measured following sample exposure.

The electrochemical behavior of the as-fabricated electrodes was initially evaluated using cyclic voltammetry (CV) in a 5 mM [Fe(CN)_6_]³ and 5 mM [Fe(CN)_6_] redox electrolyte. Scan rates between 10 and 100 mV/s were evaluated to characterize the mass-transport behavior. Electrode repeatability was evaluated using three independently fabricated electrodes from the same fabrication batch, with three consecutive measurements performed on each electrode under identical conditions. Variability was evaluated using the RSD. The inter-device RSD was calculated from the measurements obtained across the three electrodes, while scan-to-scan RSD values were calculated separately for each individual electrode. For concentration-response measurements, IL-1α was sequentially added to the same functionalized working electrode to obtain final concentrations of 25, 50, 100, and 200 pg/mL. Following each addition, the electrode was thoroughly rinsed, and the normalized ΔRct response was determined by EIS. The resulting response was plotted against the logarithm of the nominal IL-1α concentration. The limit of detection (LOD) and limit of quantification (LOQ) were calculated using the residual standard deviation of the regression (S_y/x_ = 0.1283)) and the slope of the calibration curve (b = 0.906) of the base-10 logarithmic calibration curve. The LOD and LOQ were estimated using the following equations: LOD = (3.3 × S_yx_) /b and LOQ = (10 × S_yx_)/b. Since the calibration curve was constructed using the base-10 logarithm of concentration, the calculated values were back-transformed to obtain the corresponding concentrations in pg/mL. For selectivity experiments, IL-1α was analyzed at 100 pg/mL either alone or in a cytokine mixture containing IL-1α, IL-1β, IL-6, and IL-8, each at 100 pg/mL. A corresponding mixture containing IL-1β, IL-6, and IL-8 but lacking IL-1α was used to evaluate the contribution of the non-target cytokines.

### 4.15. Statistical Analysis

Statistical analyses were performed using GraphPad Prism 10 and R. Data are presented as mean ± SD unless stated otherwise, and n refers to independent biological replicates or independently fabricated electrodes as specified in each figure legend. Two-group comparisons used paired or unpaired *t*-tests as appropriate. Multiple-group and time-course comparisons used one-or two-way analysis of variance (ANOVA) followed by Tukey’s or Dunnett’s multiple-comparison tests (on log-transformed data where applicable, e.g., CFU counts). A *p*-value < 0.05 was considered statistically significant.

## Author Contributions

Conceptualization: L.Y, D.V.C, O.P. Methodology: L.Y, D.V.C, S.X, M.H, K.L, A.Z, A.K, N.Z, M.M, C.B, A.J.M.R, O.P. Investigation: L.Y, D.V.C, S.X, O.P. Formal analysis: L.Y, D.V.C, K.L. Visualization: L.Y, D.V.C, M.H, M.M, K.L. Resources: A.Z, A.K, N.Z, M.M, C.B, A.J.M.R, O.P. Supervision: M.H., A.J.M.R, O.P. Funding acquisition: O.P. Writing—original draft: L.Y, D.V.C, O.P. Writing—review & editing: L.Y, D.V.C, S.X, M.H., K.L., M.M., A.Z., A.K, N.Z., M.M.M, C.B, A.J.M.R, O.P.

## Acknowledgements

O.P. gratefully acknowledges Karolinska Institutet (2-1534/2020), the Swedish Research Council (2022-01436) and the Bo Rydin Foundation (F04/22) for financial support. M.M. thanks Karolinska Institutet, Department of Oncology and Pathology, for financial support. AJM-R acknowledges funding from the Spanish Ministry of Science, Innovation and Universities (Agencia Estatal de Investigación, Ref. PID2023-150655NA-I00) and the Swedish Research Council for Sustainable Development (FORMAS, Ref. 2023-01097). We thank Jos P.H. Smits and Ellen H. van den Bogaard (Department of Dermatology, Radboud Research Institute for Medical Innovation, Radboud University, Nijmegen, The Netherlands) for generously providing the N/TERT-2G keratinocyte cell line on behalf of James Rheinwald (Department of Dermatology, Brigham and Women’s Hospital, Harvard Medical School, Boston, MA, USA). We also thank the Jakob Wikström group (Department of Medicine, Solna, Division of Dermatology and Venereology, Karolinska Institutet) for support with TEER measurements, and the Histological Core Facility, Biomedicum Imaging Core Facility (BIC) and Electron Microscopy Core Facility (EMil) at Karolinska Institutet for assistance with sample preparation, imaging and measurements, and research assistant Olivia Rudqvist for earlier experimental set up for biosensing.

## Ethics Statement

This study used the immortalized human keratinocyte line N/TERT-2G and previously described bacterial strains. No primary human material, identifiable human data or animal experiments were involved, and no ethical approval was therefore required.

## Data Availability Statement

The data that support the findings of this study are available from the corresponding author upon reasonable request.

## Table of Content (ToC)

A standardized 3D human epidermal infection model resolves the transition from commensal colonization to pathogen-driven barrier destruction and *identifies* IL-1α as a damage signal shared by *P. aeruginosa* and *MRSA*. The same model then supplies the biologically complex matrix in which a flexible, laser-patterned Au/polyimide immunosensor is evaluated, linking infection biology directly to animal-free bioelectronic testing.

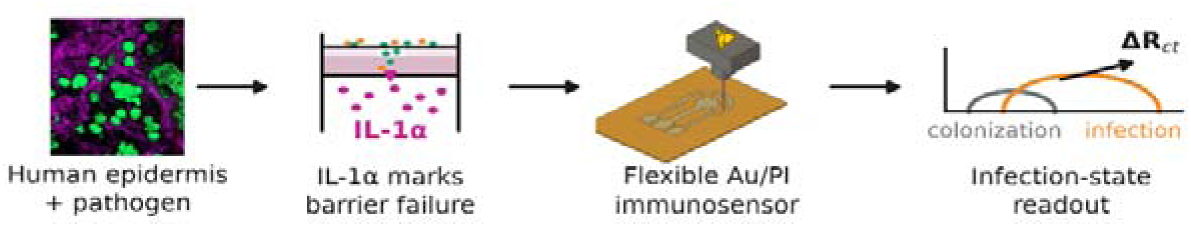

