## Supporting Information for "Profiling host-pathogen dynamic in a 3D human epidermal infection model and real-time sensing of IL-1α as a tissue-damage biomarker"

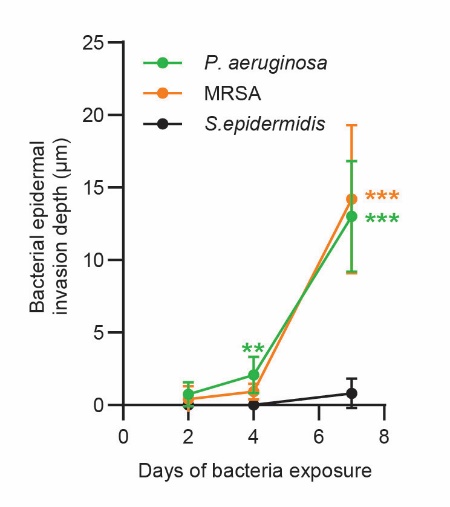

**Supplementary Figure 1. Quantitative analysis of bacterial invasion depth.** Data represent the mean ± SD of n = 5 independent human epidermal equivalent (HEE) models per group, where each replicate value represents the average invasion depth of three distinct bacterial microcolonies measured within a representative field of view from each individual model. Statistical significance at day 7 was determined by One-way ANOVA with Tukey’s multiple comparisons test (**p* < 0.05, ***p* < 0.01, ****p* < 0.001).

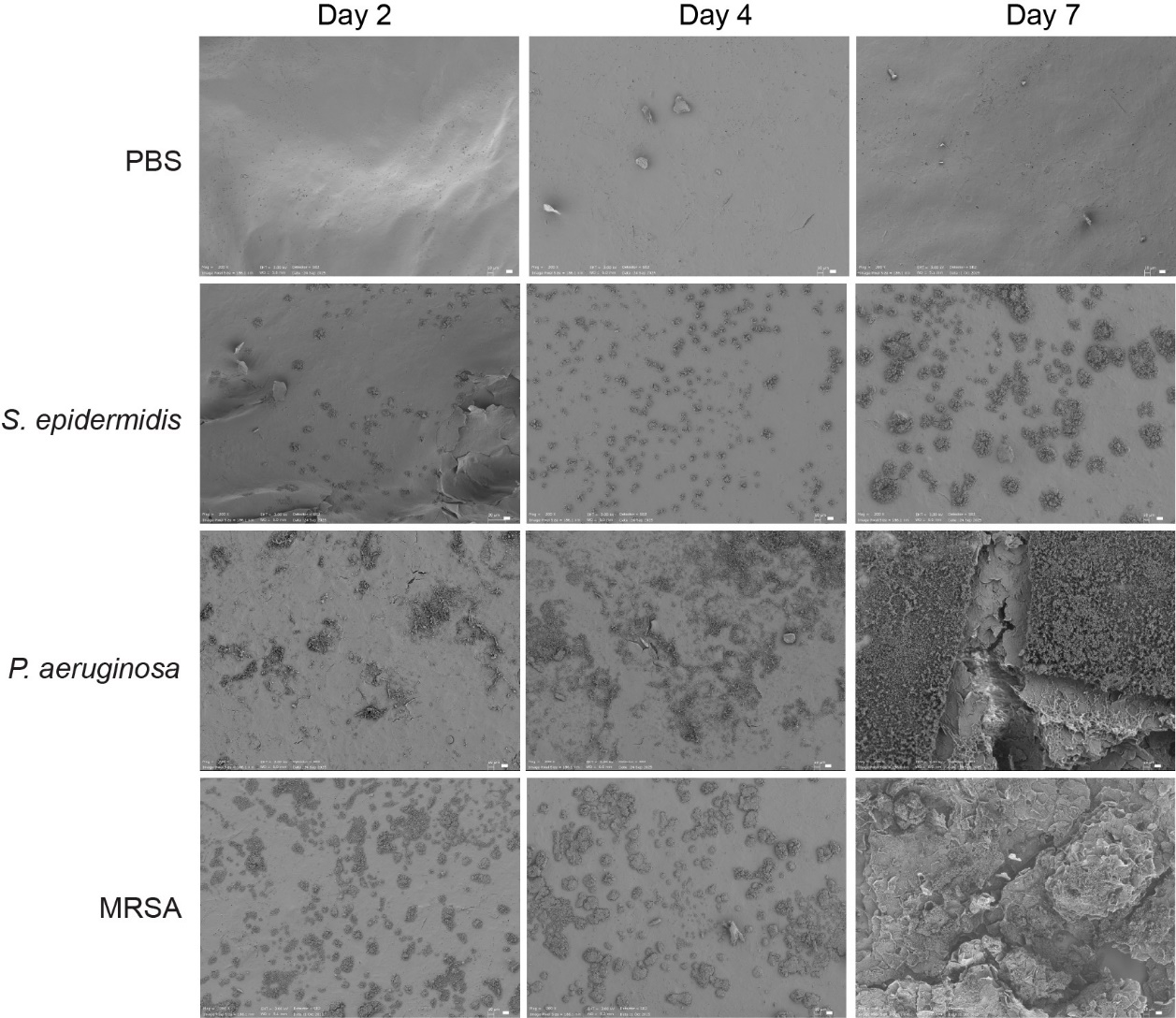

**Supplementary Figure 2. Surface topographical evolution of bacterial colonization on epidermal models.** Representative scanning electron microscopy (SEM) images from the top view of the HEE surfaces at days 2, 4, and 7. Scale bars, 10 μm.

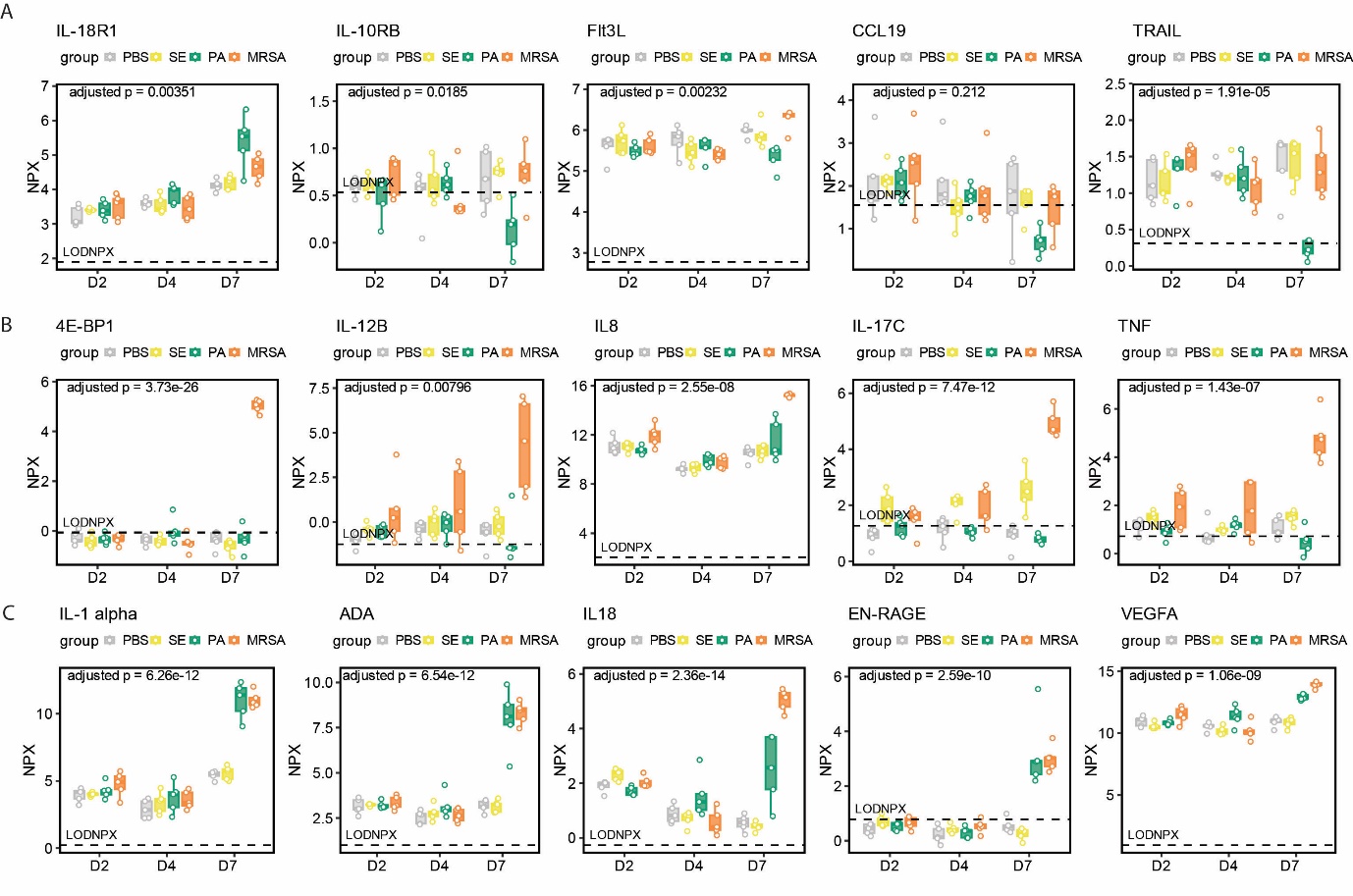

**Supplementary Figure 3.** Expression profiles of representative shared and strain-specific host proteins. Boxplots showing protein abundance (NPX) in conditioned media at days 2, 4, and 7 (D2, D4, D7) for PBS (grey), *S. epidermidis* (SE, yellow), *P. aeruginosa* (PA, green), and MRSA (orange) groups (n = 5 per group). (A) Representative proteins showing *P. aeruginosa*-specific changes. (B) Representative proteins predominantly induced by MRSA at day 7. (C) Representative proteins co-upregulated by both *P. aeruginosa* and MRSA at day 7. Horizontal dashed lines indicate the limit of detection (LOD). Adjusted p-values at the top of each plot indicate the significance of the group-by-time interaction from linear mixed-effects models.

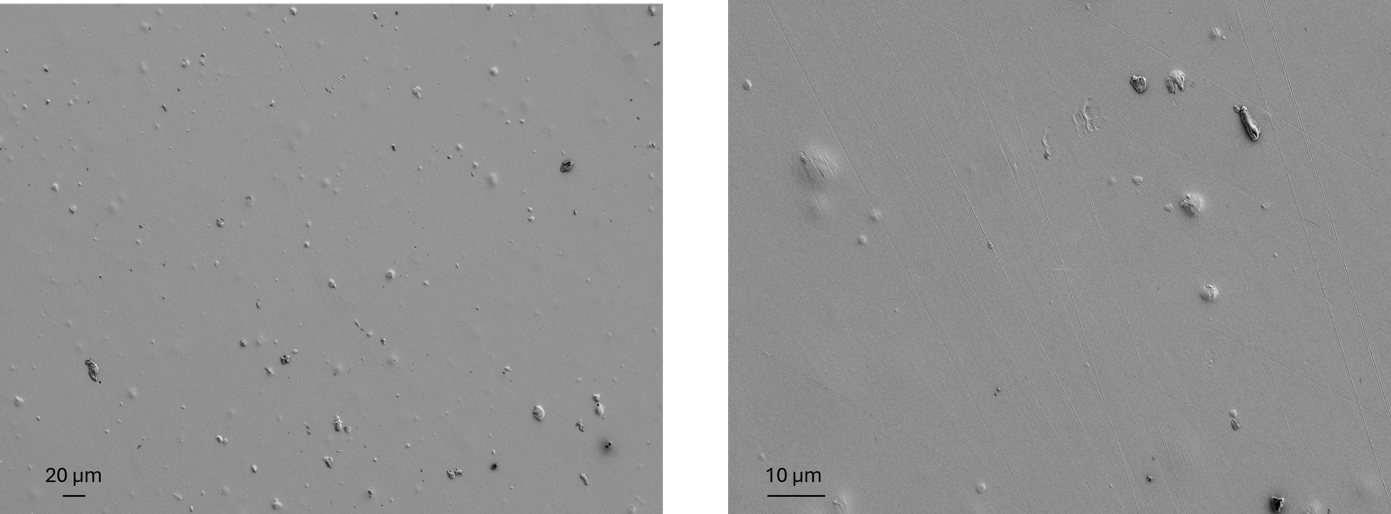

**Supplementary Figure 4.** SEM images of the gold fabricated electrodes (WE) after polyimide sheet sputtering and laser patterning geometric at different magnifications (left 200X, right: 1000 X).

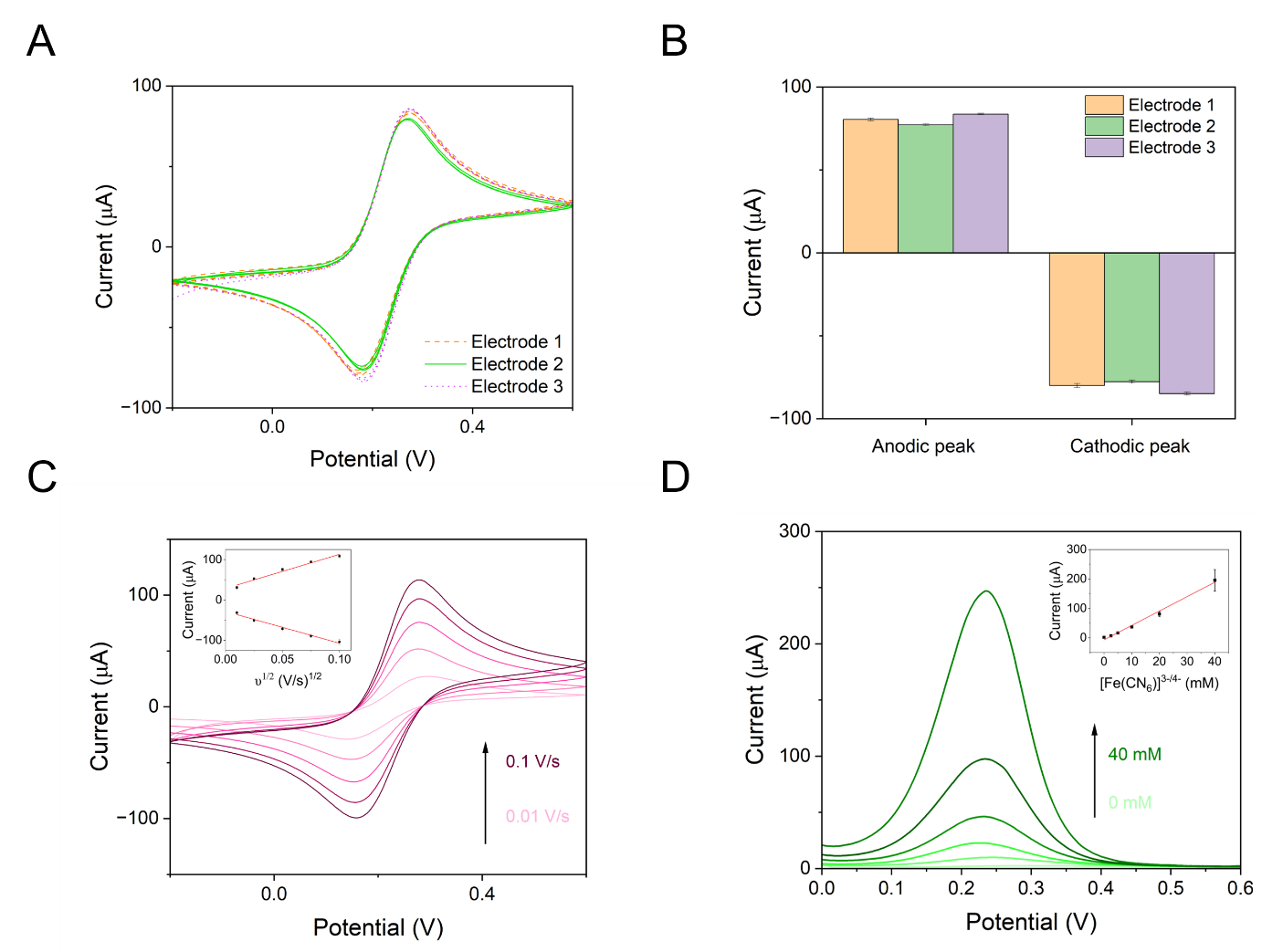

**Supplementary Figure 5. Characterization of the electrochemical performance of the fabricated Au electrodes.** (A) Reproducibility and repeatability assessed by cyclic voltammetry (CV) from three consecutive measurements using three independent electrodes in 5 mM Fe(CN)₆³⁻/⁴⁻ at a scan rate of 0.05 V/s. (B) Bar chart showing the mean anodic peak and cathodic peak current obtained from three consecutive measurements using three independent electrodes (N = 3). (C) CV curves recorded at different scan rates for evaluation of the Randles–Ševčík relationship. Inset: linear plot of the anodic and cathodic peak current versus the square root of the scan rate using three independent electrodes (N = 3). (D) Differential pulse voltammetry (DPV) curves showing the electrochemical response to different concentrations of the Fe(CN)₆³⁻/⁴⁻ redox probe. Inset: linear plot of the anodic peak current versus redox probe concentration using three independent electrodes (N = 3).
